# Adaptive evolution of carnivory in *Nepenthes* pitcher plants: a comparative transcriptomics and proteomics perspective

**DOI:** 10.1101/2023.01.07.523112

**Authors:** M Scharmann, F Metali, T U Grafe, A Widmer

## Abstract

While trait diversity associated with evolutionary radiations is easily recognized, their function and adaptive relevance often remain elusive. Here we study the evolution of carnivory genes in *Nepenthes*, an iconic radiation of carnivorous pitcher plants. We investigate 13 species chosen to represent the diversity of nutrient acquisition strategies, geography and climates covered by the genus. Using a combination of proteomics and transcriptomics, we discovered that *Nepenthes* secrete hundreds of enzymes and antimicrobial proteins into their digestive fluids. Further genes related to plant carnivory were uncovered by analyses of gene expression changes induced by experimental feeding of starved *Nepenthes* traps. Feeding status appears to affect the relative abundance of nearly 35% of all expressed genes, and may be accompanied by a strong physiological shift away from photosynthesis towards heterotrophy, proteolysis, and protein synthesis. Among the many thousand genes showing signatures of adaptive evolution in sequence or expression level within the *Nepenthes* radiation, the secreted pitcher fluid proteins and the carnivory-related genes were over-represented. A bias towards carnivory genes, and towards trap-expressed genes versus leaf-expressed genes, was also observed among the genes differing in expression in a young pair of sister species. Together, our results suggest that the molecular basis of the carnivorous syndrome was disproportionally targeted by positive selection during the *Nepenthes* radiation. This study demonstrates how the evolutionary relevance of putative key traits can be tested at the level of underlying genes.

## Introduction

Carnivorous plants have long attracted the interest of scientists [1,2]. In recent years, a solid understanding of the physiology, ecology and evolutionary origin of this enigmatic nutrient acquisition strategy has been developed [3–7]. However, we currently know very little about the subsequent diversification of carnivorous plant lineages. Carnivory has repeatedly evolved at least X times in Angiosperms, and some of these origins have established evolutionary radiations [5,8]. While it appears clear that the original gain of a carnivorous syndrome conferred selective advantages, the role that carnivory has played in subsequent radiations remains unstudied. Here we raise the question whether the carnivorous syndrome itself underwent adaptive evolution and how it has contributed to diversification, which may involve both adaptive and non-adaptive processes [9,10]. On the other hand, carnivory could be a conserved trait once established and may be tightly constrained by purifying selection, or evolve neutrally and have little effect on the subsequent evolution of a lineage.

Several frameworks exist to test whether specific traits have played important roles in evolutionary radiations. Such traits, often referred to as key adaptation traits, contribute disproportionally to functional diversity and are repeatedly targeted by diversifying selection during evolutionary radiations (e.g. flower colour and spur length in *Aquilegia*, [13,14]. A popular approach are macro-evolutionary models that aim to relate the origin of particular traits to shifts in phylogenetic diversification rates [15–17]. A novel and more direct approach applies genome-wide scans for molecular signatures of diversifying selection across radiations [18–21]. Signatures of past selection can be inferred from DNA and protein sequences by analysing ratios of non-synonymous to synonymous substitutions in codon alignments [22–24]. In addition to amino acid substitutions, regulatory changes have long been hypothesised to play important roles in adaptation [25]. Recent developments now allow better inference of gene expression evolution levels through non-parametric and model-based frameworks of quantitative trait evolution [26–29], and new empirical results have established that regulatory changes can lead to adaptation [30–32]. The contributions to adaptation of gene expression changes relative to coding sequence (structural) changes, however, remains an important question [30,33,34]. Comparative datasets of multiple species in a radiation and covering representative samples of organismal genomes or transcriptomes hold the potential to reveal the molecular basis of adaptations and to answer questions on the contributions of gene sequence and gene expression changes to adaptation.

Plant carnivory is a complex quantitative trait whose genetic basis is largely unknown. For obvious reasons, plant carnivory genes cannot directly be studied in traditional model plants such as Arabidopsis. Nevertheless, carnivory is well suited for exploration of some of the underlying genes using a combination of proteomics and transcriptomics [35]. Prey digestion in carnivorous plants is a physiological trait for which direct links can be established between the digestive enzymes, the genes encoding these enzymes, and their role in carnivory. Most available information on enzymes used by carnivorous plants to digest their prey comes from *Nepenthes* (Nepenthaceae, Caryophyllales). As early as 1874, it was noted that the fluid of *Nepenthes* pitcher plants degrades meat, egg white and cartilage [1], but digestive enzymes and their secretion were characterised only in the second half of the 20th century [3]. A major debate ensued concerning the origin of these enzymes from either plant or microbes living in the pitcher fluid (reviewed by [36]). While it has been established that microbes contribute to the enzymatic activity and nitrogen release in natural pitcher fluid [37,38], digestive enzymes are present and active even in sterile pitchers [3,39]. Evidence for plant-derived digestive enzymes was corroborated by sequencing the most abundant group of proteins found in pitcher fluids, aspartic peptidases called Nepenthesins [40], and their subsequent cloning from the plant genome [41]. The combination of proteomic and genomic techniques has since verified that *Nepenthes* secrete into their pitcher fluid a cocktail of hydrolytic and antimicrobial proteins [6,42–46], which we subsequently refer to as pitcher fluid proteins (PFPs). As they are directly degrading prey and potentially control pitcher microbial communities, PFPs are key elements of plant carnivory. At least some PFPs are not constitutively produced but are induced only after prey capture [47,48] through upregulation of gene expression in the glandular, submerged pitcher wall that is in direct physical contact with the pitcher fluid [42,49–51]. These observations suggest that sensory mechanisms detect prey and that PFP synthesis may be energetically expensive for the plant.

The genus *Nepenthes* (Nepenthaceae, Caryophyllales) is highly diverse [12,52,53], and has accumulated approximately 150 species within only c. 5 millions years [54]. Although all *Nepenthes* produce pitcher traps, there is a large diversity in morphology, biomechanics, and chemistry of the digestive fluid [12,52,53] that corresponds in part to differences in prey spectra. *Nepenthes* include insect-eating generalists, a termite specialist, a leaf-litter specialist, and several coprophagous species that attract small vertebrates and intercept their excreta [53,55]. To what extent pitcher trap diversity is also reflected at the physiological level of prey digestion remains an open question. To date, only two studies have suggested that variation in proteinase activity and PFP composition exists among *Nepenthes* species [56,57].

Here we combine proteomics and transcriptomics to investigate carnivory genes in *Nepenthes* pitcher plants. We explore the diversity of PFPs and changes in gene expression associated with feeding in the pitchers, as well as shifts in gene expression in a young species pair. We then examine signatures of natural selection on protein-coding sequences and gene expression levels in a comparative transcriptomic dataset of 13 *Nepenthes* species, selected from a broad geographic range and disparate ecology (Figure 1). We show that thousands of genes experienced positive selection on amino acid substitutions, and shifted in their expression levels during the *Nepenthes* radiation. PFPs were preferentially targeted by positive selection and experienced shifts in expression levels, while genes upregulated during feeding were overrepresented among the genes showing expression changes between recently diverged species.

**Figure 1.**
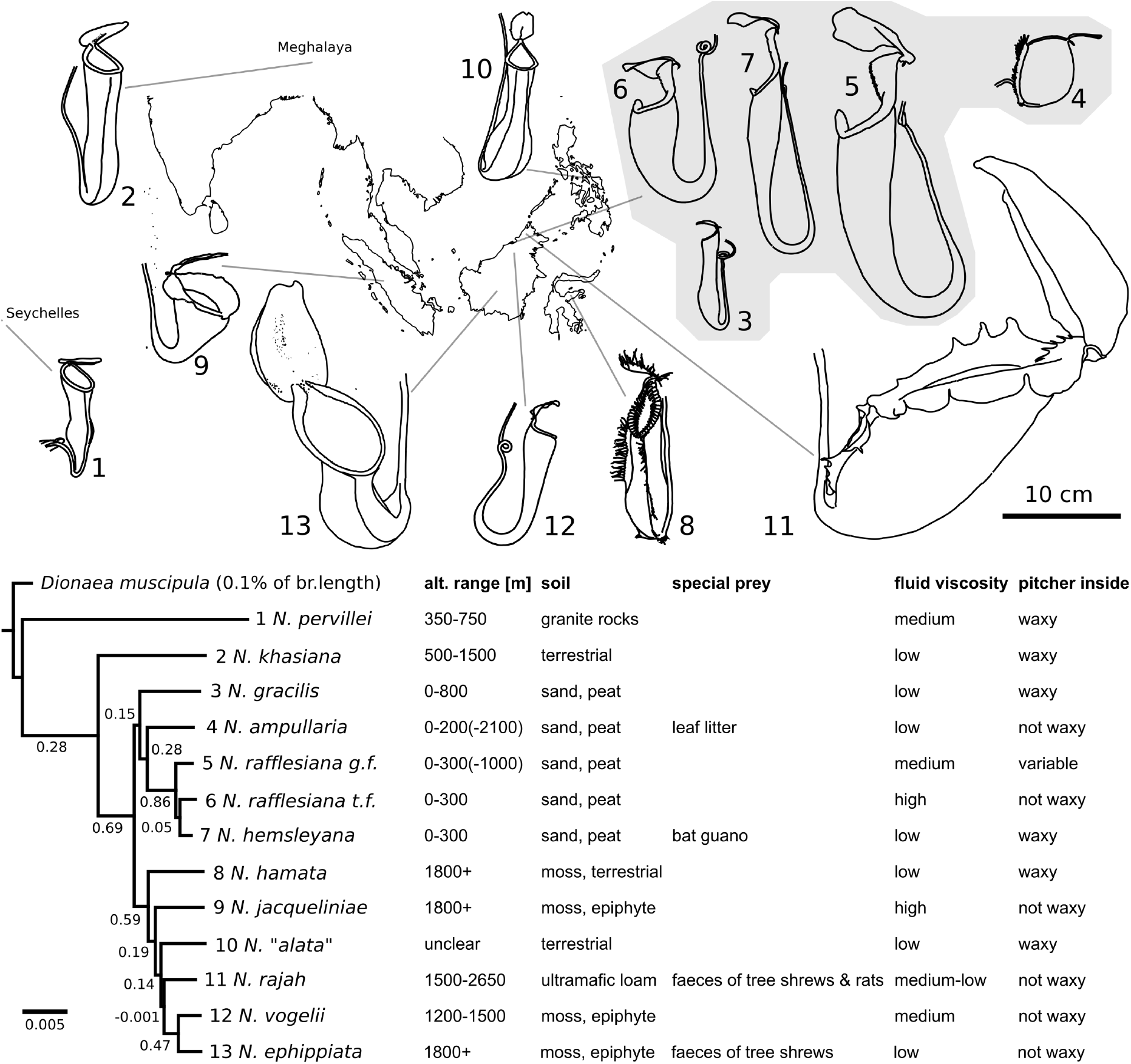
Top: Portraits of the characteristic pitcher traps produced by the 13 *Nepenthes* species examined in this study with a map of their provencance in Southeast Asia, India and the Seychelles. Bottom left: species tree estimated from 1,722 one-to-one orthologous genes for the 13 *Nepenthes* species rooted to the Venus Flytrap, *Dionaea muscipula*. Node labels indicate Internode Certainty (IC, [11]. Bottom right: major ecological and physiological traits. All *Nepenthes* prey on Arthropods, while some prefer special prey types. Data from [12] and own observations.

## Results

### The diversity of *Nepenthes* pitcher fluid proteins

We jointly analysed pitcher fluid proteomes and pitcher transcriptomes of twelve *Nepenthes* species. We found a much broader spectrum of PFPs across these species than has previously been known and that PFP composition and cumulative expression levels differ strongly among species (Figure 2). Predicted plant proteins identifed in pitcher fluids by mass spectrometry were clustered into high-order PFP “classes” with at least 40% amino acid sequence identity. We refer to proteins within these classes as isoforms. We inferred 27 PFP classes from 242 predicted protein sequences. Of these, twenty PFP classes have previously been identified, but seven classes are new, together with a large number of novel isoforms. The most diverse class were the already known Nepenthesins [40,41], with a total of 19 isoforms across species and up to 16 isoforms per species (*N. khasiana*, *N. hemsleyana*).Second in diversity were the only recently discovered proline-cleaving Neprosins [46] with up to twelve isoforms per species, and BG3-like Glucanases with up to nine isoforms per species (Figure 2 left). Nepenthesins were hitherto understood as the most abundant and important digestive enzymes in *Nepenthes*, but it appears that other PFPs occur in similar isoform diversity. Although Nepenthesins were the most highly expressed single PFP class (mRNA levels, Figure 2 right) in seven out of twelve species, they were always outnumbered by the cumulative transcript abundance of other PFP classes, except in *N. rafflesiana* t.f. The dominance of Nepenthesins among PFP transcripts was replaced by Class IV Chitinases in *N. pervillei* and *N. gracilis*, by Serine carboxypeptidase 20-like enzymes in *N. khasiana*, by Thaumatin-like proteins (cluster 2) in *N. ampullaria*, and by Class III Chitinases in *N. ephippiata* (Figure 2 right). Furthermore, the 13 species displayed a more than eleven-fold variation in cumulative PFP expression levels across all classes and isoforms. This was exemplified by the low extreme *N. ampullaria*, a detritivorous species with unusually weak fluid acidity, and the high extreme *N. vogeli*, an insectivorous species (Figure 2 right top row). Together, these results revealed a hitherto unknown PFP diversity across the genus *Nepenthes*.

**Figure 2.**
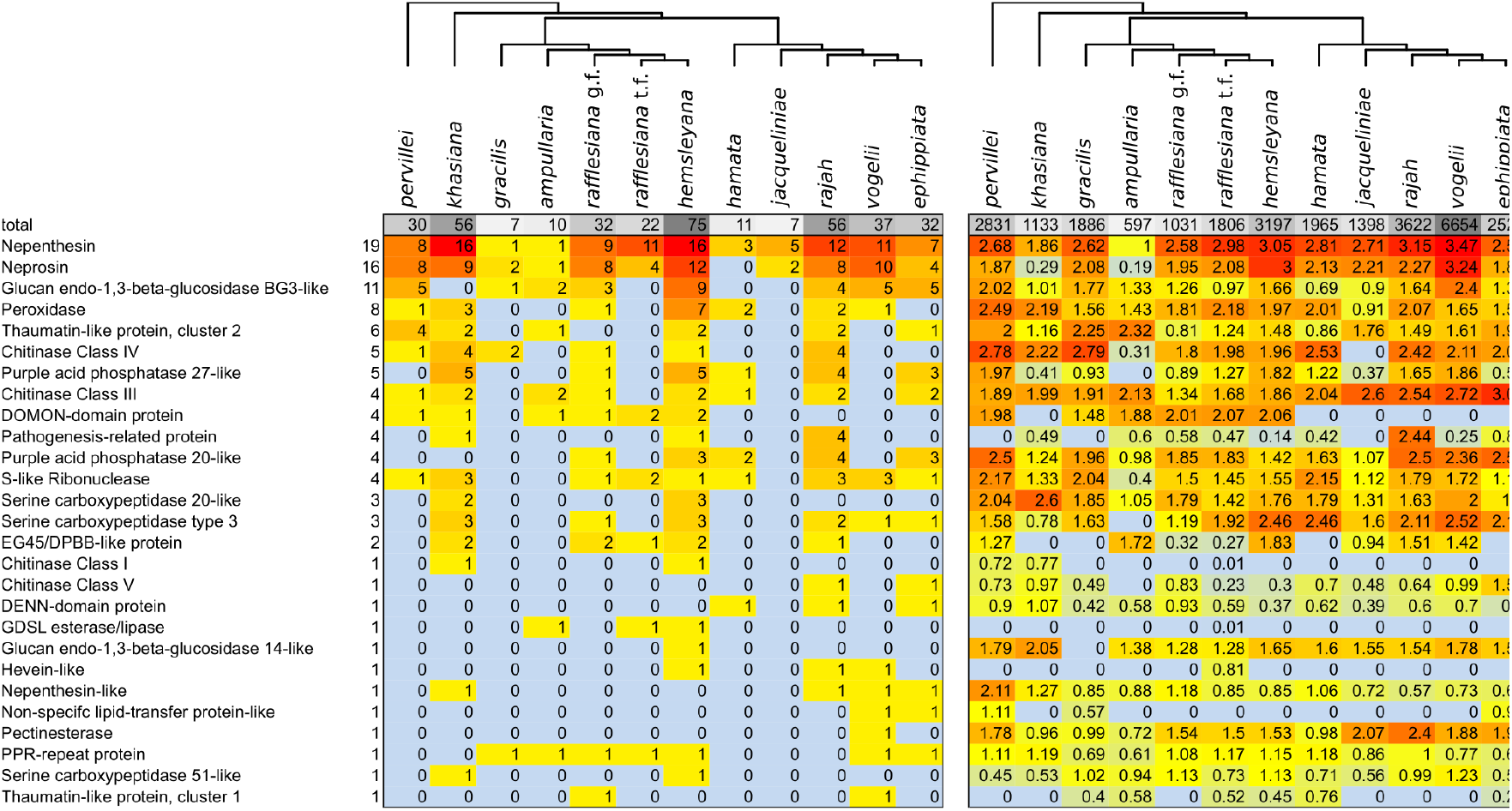
A) Plant-derived pitcher fluid protein (PFP) classes identified by mass spectrometry in pitcher fluids of 12 *Nepenthes* species seven days after elicitor treatment. Shown are numbers of distinct ‘isoforms’ per PFP class. Note that failure to detect proteins is not evidence of absence. B) Gene expression levels for the same protein classes in the glandular pitcher walls three days after feeding with *Drosophila*, measured cumulatively over all homologs present in the transcriptomes of each species. The units in the top row (total) are TMM-normalised TPM expression values, and scaled by log10(x+1) in the rest of the table. Dendrograms above tables indicate the species tree.

In addition to the numerous isoforms in 20 previously known PFP classes, we here report seven novel PFP classes. A DENN-domain protein, detected in three species, was most similar to *Arabidopsis* SCD1 (AT1G49040), which may be involved in cytokinesis but also clathrin-mediated endocytosis and reaction to pathogens [58]. A PPR-repeat protein was detected in nine samples. Members of this large protein family are known to mediate organellar RNA editing [59]. A pectine methyl esterase with strongest similarity to *Arabidopsis* PME17 (AT2G45220) was found only in *N. vogelii*. These enzymes are known to aide in pathogen defense and to modify the rheology of cell wall components [60], and it is tempting to speculate that this enzyme is involved in regulating the visco-elasticity of *Nepenthes* pitcher fluids [61,62]. Four samples yielded a Nepenthesin-like protein with only 31% amino acid identity to any canonical Nepenthesin but consistent PFAM domain structure. A Class V Chitinase (most similar to *A. thaliana* AT4G19810) was found in two species. Four samples contained Class I Chitinase proteins, which were already known to be expressed in pitcher tissues [42]. This gene family underwent sub-functionalisation into carnivory versus pathogen defence in carnivorous Caryophyllales [63], but the protein was previously not documented in *Nepenthes* digestive fluids. Two isoforms of an EG45/DPBB-like protein including signal peptides were found in six samples each. In conclusion, these proteomes and transcriptomes revealed that the secreted protein cocktails of *Nepenthes* pitcher fluids are more complex than previously known, and highly variable between species.

### Gene expression changes in feeding *Nepenthes* pitchers

Similar to an animal’s gut, the process of prey digestion in *Nepenthes* involves secreted enzymes, as well as microbes and macro-invertebrates [38], followed by absorption and assimilation of the released components. Hence, numerous physiological processes in the pitcher tissue fulfil essential supportive tasks during feeding. By comparing gene expression in fed pitchers against controls (unfed pichers) and against non-carnivorous leaves, we inferred candidate genes involved in these processes. Gene expression differed most strongly between fed pitchers and leaves, while control pitchers appeared intermediate between these conditions (Figure 3A). As expected, PFPs typically showed higher transcript levels in fed than in control pitchers, and higher levels in pitchers than in leaves (Figure 3 B). These findings support the notion that PFP production is prey-induced [47,48]. However, the observation that several PFPs appear more strongly expressed in leaves associated with fed pitchers than in the control pitchers may indicate that PFP expression between pitchers and associated leaves is not completely decoupled.

**Figure 3.**
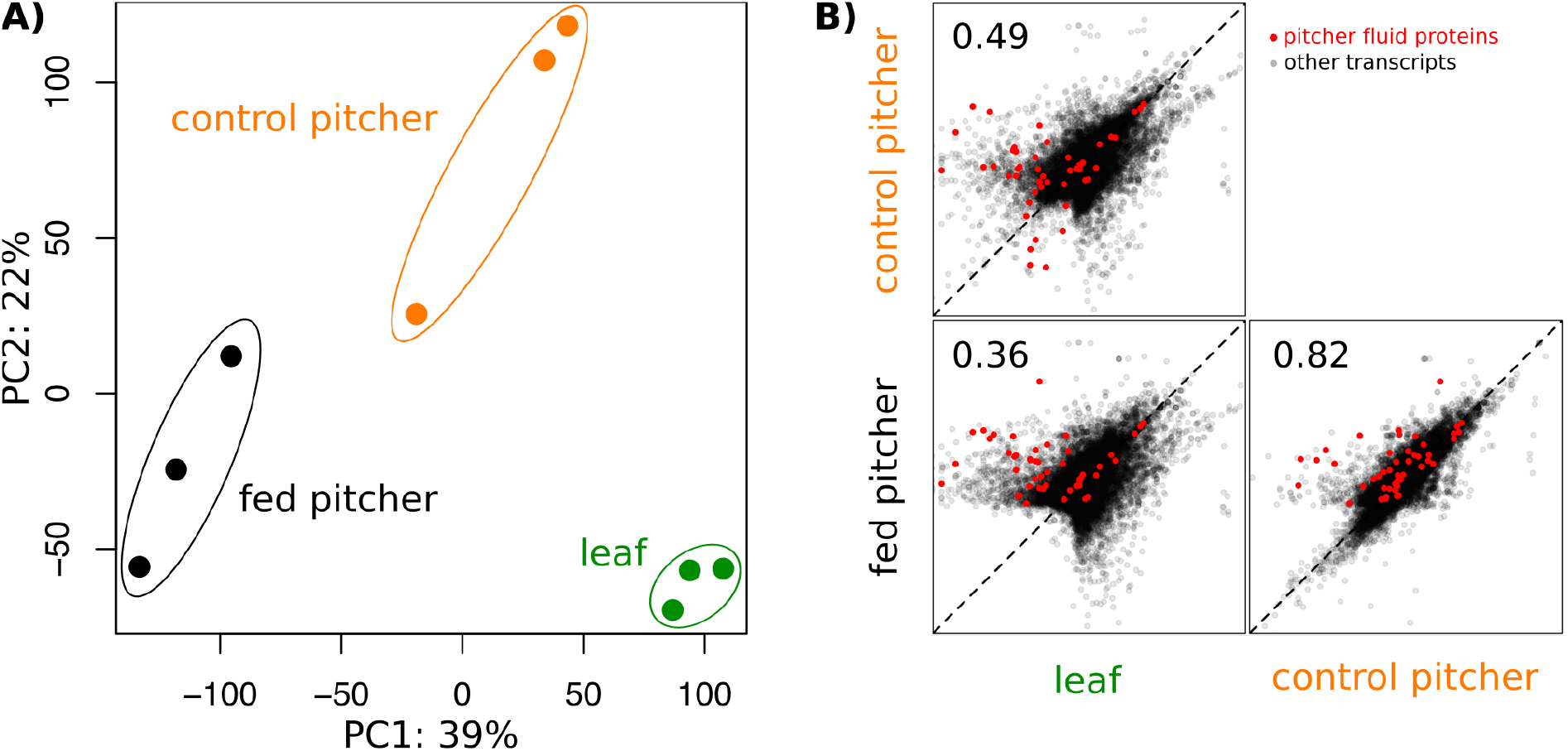
Comparison of gene expression based on 22,952 contigs between leaves, fed and control pitchers. A) Principal Component Analysis on normalised gene expression counts. B) Pairwise expression correlations between leaves, fed and control pitchers based on log2-scaled mean gene expression counts. Numbers are Pearson’s correlation coefficients; all correlations with p<2.2e-16; dotted lines indicate equal gene expression between compared tissues.

We found that 19.6% of 20,182 tested contigs were more strongly expressed in control than in fed pitchers. These were enriched for molecular functions (GO terms) related to photosynthesis, response to light stimulus, circadian clock, leaf development, and location in plastids and photosystems. In contrast, 14.7% of all contigs were more strongly expressed in fed pitchers than control pitchers and these were preferentially involved in peptide biosynthesis, translation, oxidoreduction, and associated with ribosomes. Besides the dominant up-regualtion of protein biosynthesis machinery, genes related to proteasomes, glycolysis, ATP-generation and cellular respiration were also up-regulated. The involvement of proteasomes could imply that part of the digestive process occurs intracellularly, on absorbed peptides or endocytotic vesicles [64].

Gene expression differences between fed pitchers and leaves were much stronger than between fed and control pitchers, but identified remarkably similar gene functions (Figure 3 A&B). Genes with higher transcript abundance in leaves (25.6% out of 20,595 contigs) were overall enriched for tasks in photosynthesis, carbon assimilation, chlorophyll synthesis, and chloroplastic locations. Transcripts with higher expression in pitchers (25.5%), on the other hand, were enriched for functions in aerobic respiration, nucleoside triphosphate metabolism, lipid oxidation, protein synthesis, and also for oxidoreductases and transmembrane-transporter ATPases, and preferentially located in mitochondria and ribosomes.

These results reveal that pitchers traps without prey may function rather similar to ordinary leaves. Prey capture and subsequent feeding might induce a major physiological shift from photosynthesis as the dominant process to protein biosynthesis, proteolytic activity, and respiratory energy production, i.e. a more heterotrophic state. Thus, we hypothesise that *Nepenthes* mitigates the energetic costs of carnivory by switching the central task of its pitchers from photosynthetic carbon assimilation to carnivory in response to prey capture. The ability to rapidly adjust pitcher physiology when the unpredictable demand arises may represent an adaptation that allowed *Nepenthes* to reduce energetic costs associated with maintenance of the pitcher traps.

### Gene expression differences between young sister species

We hypothesised that bias in the functions of genes differing in expression between young sister species could be indicative of directional selection on regulatory mechanisms whereas neutral divergence should lead to functionally unrelated differentially expressed genes. We investigated divergence in gene expression between the incipient sympatric sister species *N. rafflesiana* t.f. and *N. hemsleyana*. First, we fitted a two-way regression model with the factors species and tissue (fed pitchers and leaves) to the gene expression levels in the two species. Overall, differences in expression between pitcher and leaf were highly similar (correlated) between *N. rafflesiana* t.f. and *N. hemsleyana* (Figure 4 A), and multi-dimensional scaling clustered tissues before species. However, 0.8% of 25,119 tested contigs showed significant species by tissue interaction effects, i.e. the difference in expression between the tissues was species-specific (Figure 4 A red points). These genes were functionally rather heterogeneous with the few enriched GO terms dominated by regulation of the actin cytoskeleton, gluconeogenesis, flavonoid biosynthesis, and oxidoreductase activity. These genes indicate how the sister species have diverged in their regulation of genes in leaves or pitchers.

**Figure 4.**
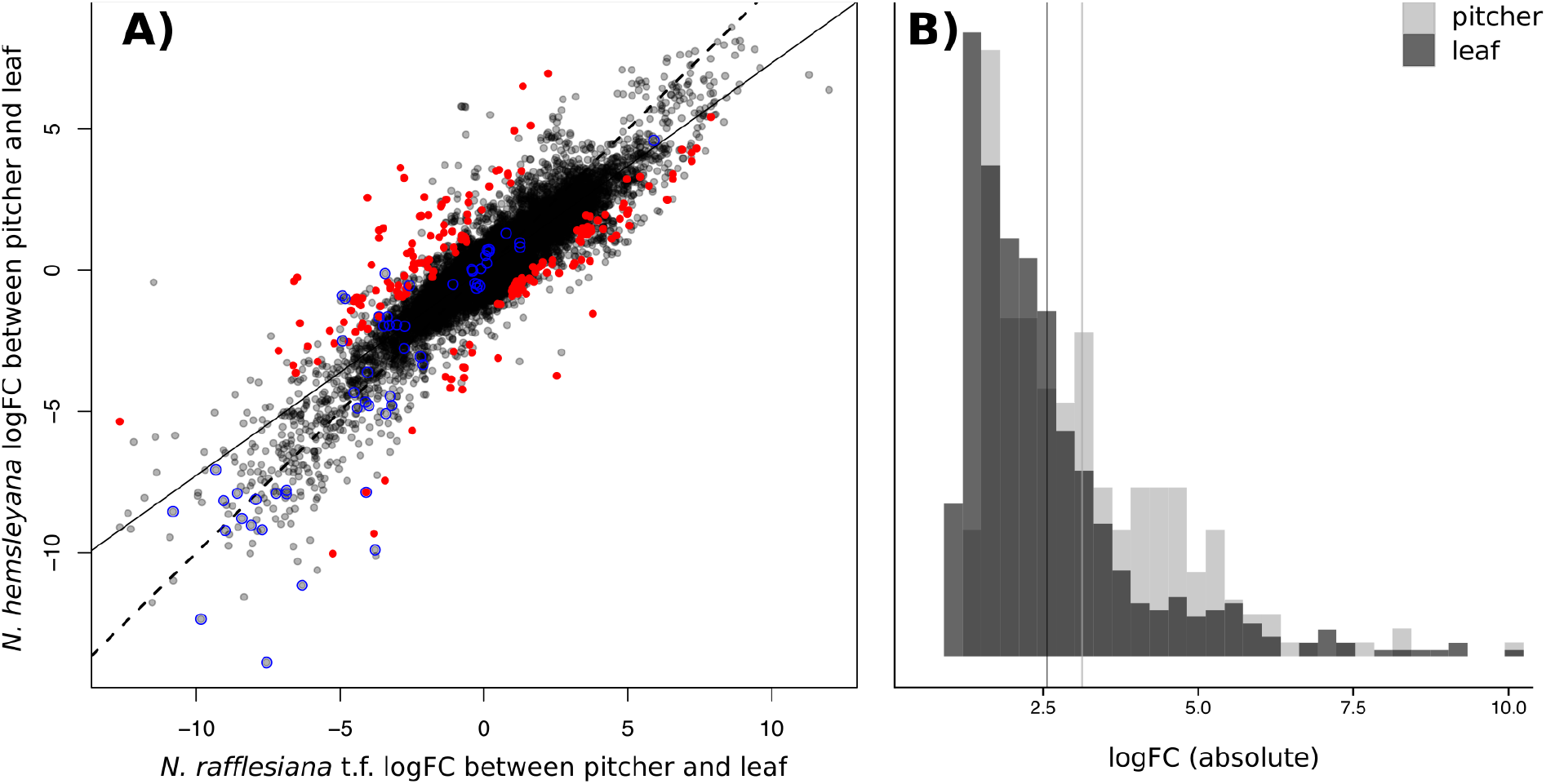
Fold-changes in expression levels between the fed pitchers and leaf tissues in the sister species *N. rafflesiana* t.f. and *N. hemsleyana*. A) correlation of fold-changes; red points: contigs showing significant tissue by species interaction, black points: interaction not significant; blue circles: PFPs; dotted line: 1:1 expectation; continuous line: linear model fit (y = 0.73 * × + 0.036, F_1,25117_ = 8.467e4, p < 2.2e-16). B) Histograms of interspecific fold-changes (log2, absolute) for DEGs between pitchers (light grey) and leaves (dark grey); distribution means indicated by vertical bars.

Second, we compared transcript levels between species for each of the two tissue types separately (but with a common normalisation). Although fewer differentially expressed genes (DEGs) were found between pitchers (1% of 25,119 contigs) than between leaves (2.1%), the magnitude of expression differences was greater in comparisons of pitchers (mean fold-change _pitchers_ = 8.75; mean fold-change _leaves_ = 5.92; permutation-p = 2e-5; Figure 4 B). This was a robust result also observed among all expressed contigs (p = 2e-5). DEGs from the pitchers of the two species were enriched for functions in secondary metabolite biosynthesis, fatty acid transport, and developmental regulators, while DEGs from the leaves were enriched for the same functions, with the notable additions of steroid synthesis and chromoplast location. The latter may correspond to leaf pigmentation, as *N. hemsleyana* developed deep red leaves and *N. rafflesiana* t.f. bright green leaves under identical culture conditions.

Pitcher–leaf differentiation in gene expression was overall lower in *N. hemsleyana* than *N. rafflesiana* t.f., as indicated by the smaller than unity slope of the correlation (Figure 4 B). This pattern was apparently dominated by a more leaf-like transcriptome of *N. hemsleyana* pitchers rather than convergence of pitchers and leaves towards an intermediate transcriptome, as the sister species were still more similar for gene expression in leaves than in pitchers (Figure 4 B). Hence, the pitchers of *N. hemsleyana* are less divergent in expression from leaves than pitchers of *N. rafflesiana* t.f., which could reflect a weaker functional specialization for carnivory or reduced investment in digestion – an interesting possibility given that *N. hemsleyana* has ‘outsourced’ some of its prey digestion to small bats which roost in its pitchers and feed the plant with guano [55,65], while its sister species is an insectivore.

Interspecific divergence in transcriptome-wide gene expression was previously reported to vary among tissues in animals [66–68], potentially as a consequence of tissuespecific selection regimes or different proportions of tissue-specific versus universally expressed genes. Our results show that gene expression between *N. rafflesiana* t.f. and *N. hemsleyana* has diverged more strongly in pitchers than in leaves. This may reflect stronger diversifying selection acting on pitchers, or instead relaxed purifying selection in the pitcher transcriptomes.

### Adaptive molecular evolution in *Nepenthes*

To understand whether adaptive evolution accompanied the diversification of *Nepenthes*, we built a comparative dataset with c. 25,000 clusters of homologous, expressed predicted proteins (irrespective of paralogy and orthology) from the *de novo* transcriptome assemblies of 13 species. We measured species-specific expression levels of these homolog clusters as the cumulative expression over all members of a homolog cluster (for 12 species, exlcuding *N. “alata”*). We hereafter refer to these homolog clusters as genes. We tested for positive selection on coding sequences, for shifts between two alternative gene expression level optima on internal branches of the phylogeny (Ornstein-Uhlenbeck process), and examined variation in expression levels based on the average of phylogenetically independent constrasts (PICs). We found that PFPs and ‘carnivory-related genes’ (972 genes that were upregulated during feeding in the pitchers and also more strongly expressed in pitchers than in leaves) were overrepresented among genes with signatures of adaptive molecular evolution.

Signatures of diversifying selection on amino acid substitions (branch-site tests) were evident in 2,688 (10.7%) out of a total of 25,079 tested genes, when enforcing a strict branch length cut-off (0.1) to remove false-positives arising from alignment errors. These genes were mildly enriched for functions (GO terms) in immune system processes and programmed cell death. Shifts between two alternative gene expression level optima on internal branches of the phylogeny were detected for 4,127 (16.3%) out of 25,258 tested genes, with no enriched GO terms. We furthermore observed that adaptations at the coding sequence were slightly depleted among genes showing expression level shifts (1.5% out of 25,075 genes, expected 1.8%, χ^2^_df=1_=8.5, p<0.01), indicating that positive selection in *Nepenthes* does typically not act jointly on regulatory and structural variation. Strong functional bias was evident among the genes in the top 5% of the transcriptome-wide distribution of gene expression variation (mean PICs): they are involved preferentially in translation, peptide synthesis, nitrogen metabolism, photosynthesis, and aerobic respiration, and localised in ribosomes, chloroplasts and mitochondria. Interestingly, these GO terms were very similar to those retrieved for gene expression changes induced by feeding.

We then tested our main hypothesis that genes underlying carnivory show accelerated molecular evolution in *Nepenthes*. We could match 49 genes of the comparative dataset to PFPs, 15 of which showed signatures of positive selection at the coding sequence level. This is an approximately threefold over-representation of this functional class among all genes exhibiting signatures of positive selection at the coding sequence (Table 1, first column). PFPs were furthermore over-represented among genes displaying shifts in expression level across the phylogeny, had elevated expression variation between species, and were overrepresented among the genes diverging in expression in a young species pair (Table 1).

**Table 1.** Interrelations between patterns of molecular evolution and the genetic basis of carnivory in *Nepenthes* (PFPs and carnivory-related genes), as tested in 2×2 contingency tables and permutation tests, respectively. Branch-site tests for positive selection on amino acid substitutions were conducted across 13 species. Gene expression levels were known for 12 species and tested for shifts in optimal expression using models of quantitative trait evolution (Ornstein-Uhlenbeck process), as well as non-parametrically compared using phylogenetically independent contrasts (PICs; permutation tests on the difference in means). Speciation-related genes are defined here as genes with differential expression between *N. rafflesiana* t.f. and *N. hemsleyana*, in any of pitchers, leaves or with an interaction tissue by species (see previous section).

|  | branch-site test |  | Ornstein-Uhlenbeck shift of optimal expression level |  | expression level mean of PICs |  | speciation-related |  |
| --- | --- | --- | --- | --- | --- | --- | --- | --- |
|  | significant | ns | shift | no shift | N | mean | yes | no |
| **PFPs** | 15 | 34 | 14 | 35 | 49 | 4.8 | 5 | 24 |
| **non-PFPs** | 2673 | 22357 | 4113 | 21096 | 25209 | 3.1 | 253 | 8014 |
| $\chi^2$ | 18.3 | | 4.5 | | na | | 14.9 | |
| p-value | 2e-5 |  | 0.03 |  | 2e-5 |  | 0.0001 |  |
| **carnivory-related** | 138 | 833 | 174 | 798 | 972 | 4.6 | 40 | 932 |
| **not carnivory-related** | 908 | 4620 | 886 | 4669 | 5555 | 4 | 112 | 5443 |
| $\chi^2$ | 2.8 | | 2.2 | | na | | 15.1 | |
| p-value | 0.09 |  | 0.14 |  | 2e-5 |  | 0.0001 |  |

We identified as ‘carnivory-related genes’ those 972 genes with higher expression in fed pitchers compared to control pitchers or leaves in *N. rafflesiana* t.f. These genes were between species on average more variable in their expression (mean PICs) than other genes (Table 1), which may be a consequence of elevated phenotypic plasticity, relaxed stabilising selection, or diversifying selection. Interestingly, though, genes under positive selection or displaying shifts in expression were not overrepresented among carnivory-related genes (Table 1). We hypothesize that they encompass a high proportion of genes involved in fundamental cellular processes such as respiration and translation, and that these genes are under purifying selection. However, carnivory-related genes included three PFPs (a Nepenthesin, a purple acid phosphatase 20-like protein and a peroxidase) and were overrepresented among the genes that are differentially expressed between young sister species (Table 1), potentially indicating that prey digestion is a quantitative trait that quickly diverges between *Nepenthes* species with different prey spectra (insects versus bat guano).

## Discussion

Our combined proteomic and transcriptomic analysis of 13 *Nepenthes* species reveals that carnivory is a complex trait based on many genes, and has evolved substantial variation between species in both gene expression, as well as protein composition and abundance. We uncovered signatures of adaptive evolution in the coding sequences and expression levels of genes underlying carnivory. While numerous other genes not directly related to feeding and carnivory also exhibited adaptive divergence at the coding sequence or changed their expression levels during the *Nepenthes* radiation, those linked to feeding were in almost all our tests overrepresented – between incipient species as well as in the phylogenetic comparative dataset. These findings support the notion that plant carnivory underwent adaptive evolution subsequent to its origin and contributed to the adaptive divergence between species as a key adaptation trait in the radiation of *Nepenthes*.

Our study is among the first to test at the molecular level the hypothesis of the existence of a key adaptation trait in an evolutionary radiation. We base our argument for adaptive evolution on functional bias in genes showing elevated sequence or expression level divergence rather than rates of change or speciation. Neutral diversification is not expected to affect functionally similar or interacting genes, but should instead result in elevated divergence of genes with unrelated functions. A recent study of heavy-metal tolerance in *Arabidopsis halleri* [69] concluded that the genetic basis of complex organism-level functions conferring ecological adaptation can indeed consistently display molecular signatures of positive selection, in this case in expression. Similarly, transcriptome-wide tests for convergent coding sequence adaptations discovered novel genes involved in C4 photosynthesis in grasses [70], and comparative transcriptomics of the *Lupinus* radiation in the Americas revealed positive associations between phylogenetic diversification rate and coding sequence adaptations, as well as shifts in gene expression [21]. This study, however, could not directly test whether life history (perenniality), the suspected key adaptive trait in this radiation, was indeed the target of diversifying selection during this radiation, because genes controlling this complex trait are largely unknown. To reliably infer the role of selection in driving adaptive divergence of key ecological or morphological traits thus still requires knowledge of the genes underlying a given trait.

In comparative studies of evolutionary radiations, tests of selection in both coding sequence and gene expression levels are still rarely applied on a genome-wide or transcriptome-wide scale (but see e.g. [18]). Our study exemplifies the strength of this approach if genes can be linked to the trait of interest. It allows discovering adaptations without bias from pre-conceived hypotheses by contrasting signatures of molecular evolution between different groups of genes in the same species, and may discover functional bias in genes that have experienced similar selection regimes [71]. We expect that the reduced costs of sequencing, together with innovative statistical advances [27,29], will stimulate wider applications of these pioneering comparative transcriptomic approaches [21,28,68,72]. In contrast to many other studies focusing exclusively on signatures of selection in coding sequences, we also addressed gene expression evolution and found that greater expression change higher incidence of shifting expression optima strongly associated with a presumed adaptive trait. This finding supports the long-standing hypothesis that regulatory changes may drive adaptations [34].

We uncovered a strong transcriptomic reaction of *Nepenthe*s traps to feeding. The trend away from photosynthesis towards proteolysis, protein synthesis and respiration is mirrored in the Venus flytrap (*Dionaea muscipula*, [73], the only other carnivorous plant in which the transcriptomics of feeding has been studied to date. Experiments are required to clarify to which extent these transcriptional changes translate to physiology. Of particular interest is the net energy balance of *Nepenthes* pitchers, and whether the increased respiratory activity during feeding sources metabolites from internal storage, normal photosynthetic leaves, or from the prey, fuelling their own digestion. Such tests might resolve whether plant carnivory, thus far understood as a mineral acquisition strategy, involves partial heterotrophy. Recent experiments of gas-exchange and feeding with ^13^C-labelled prey indicate that *Dionaea* may indeed fuel its respiratory energy production with organic carbon from prey [74]. Such studies give fascinating new insights into general carnivorous plant physiology, and could in the future include comparative experiments to understand how carnivorous plant physiology has adapted to different prey types and environments.

Finally, another highly important question posed by our results concerns the role of presumed adaptions in carnivory-related genes and PFPs in speciation. Are these candidate adaptive genes directly involved and possibly contributing to the evolution of reproductive isolation, e.g. by tapping new types of prey or better control of micro-organisms, or do they represent secondary adaptations that follow suit other major evolutionary changes? We speculate that both of these scenarios occurred among the 13 ecologically diverse *Nepenthes* studied here (Figure 1). Genes that exhibit adaptive evolution in comparative genome- or transcriptome-wide datasets at the level of evolutionary radiations should be tested for divergence between incipient species and populations. Such dual-level studies may reveal the overlap between the targets of positive selection over short and long evolutionary timescales, on both structural and regulatory variation, and improve our understanding of the predictability and repeatability of adaptive divergence.

## Methods

### Plants

Lowland species including those used for differential expression analyses (*N. ampullaria*, *N. hemsleyana*, *N. rafflesiana* t.f., *N. rafflesiana* g.f., *N. gracilis*) were cultivated in a growth chamber at a constant temperature of 27°C, 90-100% relative humidity, 12 hours illumination, and in *Sphagnum-perlite* substrate with tray watering. Plants were regularly fertilised and treated with pesticides but not for several weeks prior and during experiments. All other *Nepenthes* were grown under variable but generally cooler conditions, either in greenhouses under natural lights or in terraria under fluorescent bulbs.

Raw data for greenhouse-grown *N. alata* was previously published [6]. These plants belong to a single clone originating in the Philippines, but no further details on the provenance are known (pers. comm. K. Fukushima, Sh. Yamada). According to current taxonomy [75] this glabrous specimen possibly represents *N. graciliflora* rather than *N. alata*.

### RNA treatment and sampling

For analysis of expression changes by feeding, we fed separate pitchers of three different genotypes of *N. rafflesiana* t.f. either with powdered and freeze-dried *Drosophila* flies (10 mg/ml of digestive fluid, “fed”), and different pitchers of the same individuals with demineralised water (“control” / “unfed”). These treatmens were performed on mature pitchers (minimum 7d after opening of the lid) that were kept free of insects and semi-sterile by plugging the orifice with cotton whool upon opening. Treatments on the same plants were not performed simultaneously but separated by several weeks until new pitchers had grown, to avoid any possible systemic responses of feeding [76]. After 72 hours of exposure to the treatments, the submerged part of the glandular pitcher wall (directly in contact with digestive fluid) was harvested, the fluid discarded and tissue surfaces washed in 70% ethanol before flash-freezing in liquid nitrogen. The leaf tissue was a c. 4 cm^2^ part of the leaf from which the “fed” pitchers grew, harvested simultaneously and identically to pitchers.

To test gene expression differences between young species, we used three different genotypes of *N. hemsleyana* (the sympatric sister species of *N. rafflesiana t.f*., see Chapter I of this thesis), on which we performed the “fed” treatment as above and also harvested their leaves. To form the comparative dataset, ten further species with a single sample each were likewise subject to the “fed” treatment as above.

Extensive testing unfolded that the best quality and quantity of total RNA from *Nepenthes* was obtained by the Total RNA Mini Kit (Plant) (Geneaid Biotech Ltd, New Taipei City, Taiwan) with the “PRB” lysis buffer. Tissues were ground to a fine powder in liquid nitrogen by mortar and pistil and subsequent (still frozen) milling with steel beads on a shaker mill (various). To increase RNA yield from a limited amount of tissue, lysation was conducted twice and the volumes united. A DNAse digest was performed after elution, and RNA extracts kept frozen until further use.

### RNA-seq libraries and sequencing

Sequencing libraries were prepared from 0.5 μg total RNA using the NEBNext Ultra Directional RNA Library Prep Kit for Illumina (New England Biolabs, Ipswich, MA, USA), according to the manufacturer’s protocol. We sequenced two lanes with ten resp. 15 libraries each on the Illumia HiSeq 2500 for 126 bp paired-end reads at the Functional Genomics Center Zurich.

### Transcriptome assemblies

In addition to the transcriptomes of 12 species sequenced for this study, we downloaded RNA-seq raw reads for *N. “alata”* [6] from NCBI Sequence Read Archive (seven 101bp paired-end Illumina libraries, accession numbers DRR051743-DRR051749). Transcriptomes were *de novo* assembled for each species separately using Trinity [77]. The *Dionaea* transcriptome raw assembly v1.03 was downloaded from http://tbro.carnivorom.com [73]. Coding sequences and peptides were predicted from raw transcriptome assemblies with TransDecoder.LongOrfs v3.0.0 and TransDecoder.Predict [77]. Predicted proteins were discarded if lacking similarity to known plant genes (blastp e-value <= 1e-5; custom database of all proteins of all Eudicotyledonae with sequenced genomes, NCBI Genbank accessed 6 June 2016, collapsed at 70% identity with CD-HIT, [78]).

### Homology and orthology inference

We inferred orthology of predicted *Nepenthes* and *Dionaea* genes following the tree-based pipeline of [79] with slight modifications. In brief, DNA sequences were all-versus-all BLAST compared, and a table of reciprocal hit fractions (coverage percentage for either of two sequences) was constructed with cutoff value 0.5. Clusters were built using mcl [80], with parameters -I 1.4 and e-value threshold 1e-5. Clustered sequences were then aligned based on their amino acid sequence (MAFFT v7, [81] and maximum-likelihood phylogenetic trees [82] estimated on back-translated DNA sequences. To establish finer sequence clusters, trees were pruned (maximum 0.1 expected substitutions) and split based on a maximum length for internal branches of 0.05 expected substitution, which clearly excludes any gene duplications older than the *Nepenthes* crown (at max. 30 million years, assuming a high substitution rate of 1.67*10^-8^ per site per year). *Dionaea* was exempted from branch-length pruning because it consistently displayed much higher distances than within *Nepenthes*. Sequences from these clusters were submitted to the same procedure a second time, and the final clusters were considered homologs. Homologs with at most one sequence per taxon and at least five taxa were retained as 1-to-1 orthologs to build a species tree. These and additional clusters of homologs with at least four sequences, irrespective orthology or paralogy, were used to scan for selection at the coding sequence and expression levels (homolog clusters or ‘genes’). Inclusion of potential paralogs more than doubled the number of genes available for analysis, and is justified because adaptations are expected in both orthologs and paralogs (e.g. copy number variation, sub- and neo-functionalisation).

### Transcriptome and homolog cluster annotations

Predicted proteins of the *N. rafflesiana* t.f transcriptome with similarity to plant genes and majority-consensus peptide sequences of 1-to-1 orthologs were compared to UniProtKB/TrEMBL (blastp, e-value <= 1e-5, version 28 June 2016). TrEMBL is taxonomically much more diverse than UniProtKB/Swiss-Prot, which was insufficient for taxonomic classification of sequences, and computationally more feasible than NCBI nr. Proteins were discarded whose best TrEMBL hit was not to any genus of Embryophyta (NCBI Taxonomy, Entrez query “Embryophyta[subtree] AND genus[rank]”, accessed 19 February 2017). This step identified large numbers of fungal and bacterial sequences, which were expected given non-sterile plant cultivation and the pitchers’ microbiome blooming on *Drosophila* powder. Remaining proteins were compared to UniProtKB/Swiss-Prot (blastp, evalue<=1e-5, version 19 February 2017). GO terms for hits to Swiss-Prot were extracted from the UniProtKB/Swiss-Prot text database (uniprot_sprot.dat, version 19 February 2017)

Signal peptides and transmembrane domains were predicted with Phobius 1.01 [83] and SignalP 4.1 [84].

The same procedure was applied to homolog clusters, represented by the longest sequence in the cluster.

### Species tree estimation

We used 1-to-1 orthologous sequences from 13 *Nepenthes* and *Dionaea* to estimate a common species tree to be used in molecular evolution tests. There were 1,722 orthologs present in each of the 14 taxa, yielding a concatenated matrix of 2,480,430 DNA sites, with 99% occupancy. We estimated a maximum-likelihood tree on the partitioned alignment using RAxML v8.2.4 [82] with the GTRCAT substition model. The final tree was optimised and evaluated using the Shimodaira-Hasegawa-like approximate likelihood ratio test (SH-like aLRT, RAxML -f J). Full support for all nodes was indicated. As a more informative indicator of phylogenetic conflict among loci, we calculated the “internode certainty” (IC), which for a given internode quantifies its frequency in a set of gene trees, jointly with the frequency of the most common conflicting bipartition [11]. To this end, separate gene trees were estimated for each partition (RAxML, GTRCAT) and IC annotated on the overall tree based on this collection of gene trees (RAxML -f i).

### Proteomics treatment and sampling

The digestive process was induced in *Nepenthes* pitchers by known elicitors of the secretion of proteins and secondary metabolites [42,47,85], and minerals known to be absorbed by the traps of *Nepenthes* [86,87] or other carnivorous plants [88]. Digestive fluids were supplemented with an elicitor solution, and contained at the beginning of experimental treatment 0.3 mg/ml colloidal chitin (product no. C9752, Sigma-Aldrich, Buchs, Switzerland; pulverised), 100 μM jasmonic acid (product no. 14631, Sigma-Aldrich, Buchs, Switzerland), 15 mM NH_4_NO, 1.15 mM KH_2_PO_4_, 0.7 mM NaCl, 0.35 mM MgSO_4_, and 0.35 mM CaCl. This mineral stoichiometry follows approximately that of typical terrestrial insects [89,90], while the concentrations are based on the observation of up to 15 mM NH_4_^+^ during prey digestion in *N. bicalcarata* pitchers [91], and presented no unusual osmotic stress as they remained well below total solute concentrations within typical plants cells (200-800 mM, [92]).

At least seven days after opening and blocking pitcher orifices with cotton wool, digestive fluids in pitchers were topped up with ddH_2_0 to a natural fluid level, volumes were measured using syringes, and mixed with 10% 10x elicitor stock solution. Syringe tips were guarded with soft silicone tubes to avoid tissue damage. Preliminary experiments showed that elicitor addition did not harm pitchers, but within 48 h caused a drop from pH 5-6 to pH 1-3, and a slight increase in solute protein. Digestive fluids were harvested 7-8 days after treatment and sterile filtrated (Whatman syringe filter 0.2 μm, FP 30/0.2 CA-S, GE Healthcare), then stored at −20°C until further use. Before mass spectrometry, digestive fluids were evaporated under vacuum at room temperature, and 10-100 μg protein (Qubit Assay) were aliquoted and completely dried. The extreme visco-elasticity of some digestive fluids (e.g. *N. rafflesiana* t.f., *N. jacqueliniae*) was overcome by dissolving the dried residues in 30-50 μl 1M HCl and incubation for 7-15 min at 95°C (acid hydrolysis of polysaccharides).

### Protein identification by mass spectrometry

Proteins in digestive fluids were identified using gel-free shotgun LC-MS/MS analysis in combination with *Nepenthes* transcriptome sequences as reference database. Dried or concentrated digestive fluids were dissolved in buffer (10 mM Tris, 2 mM CaCl2, pH 8.2), proteins were precipitated with TCA (10% final concentration), and the pellets were washed twice with cold acetone. Dry pellets were dissolved in 45 μl buffer (as above) and 5 μl trypsin (100 ng/μl in 10 mM HCl). Digestion was carried out either in a microwave instrument (Discover System, CEM) for 30 min at 5 W and 60°C or overnight at 37°C.

For LC-MS/MS analysis, samples were dissolved in 0.1% formic acid (Romil) and an aliquot of 5-10% was analysed on a nanoAcquity UPLC (Waters Inc.) connected to a Q Exactive mass spectrometer (Thermo Scientific) equipped with a Digital PicoView source (New Objective). Peptides were trapped on a Symmetry C18 trap column (5 μm, 180 μm × 20 mm, Waters Inc.) and separated on a BEH300 C18 column (1.7 μm, 75 μm × 150 m, Waters Inc.) at a flow rate of 250 nl/min using a gradient from 1% solvent B (0.1% formic acid in acetonitrile, Romil)/99% solvent A (0.1% formic acid in water, Romil) to 40% solvent B/60% solvent A within 90 min. Mass spectrometer settings were: Data dependent analysis, precursor scan range 350-1500 m/z, resolution 70k, maximum injection time 100 ms, threshold 3e6; fragment ion scan range 200-2000 m/z, resolution 35k, maximum injection time 120 ms, threshold 1e5.

Tandem mass spectra were converted to Mascot generic format using proteowizard v.3.0.5759 with vendor peak picking option for MS/MS and deisotoped and deconvoluted using the H-Scorer script [93], as automatised in FCC [94]. Mascot (Matrix Science, London, UK; version 2.5.1.3) was used to search peak data against a custom protein database assuming non-specific enzymatic digestion, because some *Nepenthes* fluid proteins can lysate others [95]. Our database (2,803,176 entries) combined UniProtKB/Swiss-Prot with all (unfiltered) proteins predicted by TransDecoder from the transcriptomes of all twelve *Nepenthes* species we sequenced. Mascot was searched with a fragment ion mass tolerance of 0.03 Da, a parent ion tolerance of 10.0 PPM, and oxidation of methionine specified as a variable modification.

Scaffold (version 4.6.2, Proteome Software Inc., Portland, OR) was used to validate MS/MS based peptide and protein identifications. Peptide identifications were accepted if they could be established at greater than 95,0% probability by the Scaffold Local FDR algorithm. Protein identifications were accepted if they could be established at greater than 95,0% probability (Protein Prophet algorithm[96]) and contained at least one identified peptide. If multiple database proteins were not distinguished by MS/MS peptides, they were united as “protein groups” to satisfy parsimony.

### Annotation of pitcher fluid proteins

Predicted proteins from transcriptomes that were detected in digestive fluids were scanned for PFAM domains using hmmer 3.1b1 [97] with an e-value threshold of 1e-10. Signal peptides and transmembrane domains were predicted with SignalP v.4.1 [84] and Phobius v.1.01 [83]. Using BLAST [98] and e-value thresholds of 1e-5, sequences were compared to NCBI Genbank nr (version 31 March 2016), *Arabidopsis thaliana* proteins in UniProtKB (version 3 April 2016), and a custom database of previously reported *Nepenthes* digestive fluid proteins. This database of “reference pitcher fluid proteins” was established by searching for all nucleotide and protein sequences for *Nepenthes* on NCBI Genbank (accessed 10 October 2016), and exclusion of any orthogroups without evidence from digestive fluid proteomes by referring to the cited literature. Sequences from the more recent studies by [46] and [6] were also added. Sequences were flagged as “microbial” if the header or description of an NCBI Genbank nr hit did not contain any genus of Tracheophyta (downloaded from NCBI taxonomy, 25 March 2016).

For “protein groups”, each sequence was separately annotated, but annotations were collapsed into a single representative “group annotation”, if any of the following conditions were met: (1) any of the sequences were previously reported from *Nepenthes*, (2) all sequences contained the same PFAM domains, (3) all sequences matched the same *Arabidopsis* sequence, (4) all sequences had at least one PFAM domain in common. Otherwise, all of the dissimilar annotations were reported. Finally, a table was generated listing all identified protein groups with their annotations, and all analysed *Nepenthes* pitcher fluids, with their total unique peptide counts per protein group.

### Expression levels of homolog clusters

We estimated expression levels for homolog clusters and PFP families across species by mapping sequence reads back to the species-specific denovo reference assembly, using BWA-MEM [99] with default settings. Non-primary (redundant) alignments were excluded using samtools (view -F 0×0100). Read counts were extracted with samtools idxstats and converted to tags per million (TPM[100]) by a custom R function, normalising for library size and transcript length. TPM were averaged for species with multiple replicates (*N. hemsleyana*, *N. rafflesiana* t.f.). Species lacking homologs in their transcriptome assembly were assigned the expression level zero, reflecting that transcript abundance was below a detectable level. TPM were then summed within samples over all members of homolog clusters (see homology inference) to achieve a cumulative expression level. TPM values of PFP classes and homolog clusters were then normalised across samples using the TMM algorithm in edgeR [101].

### Tests for diversifying selection on homolog cluster expression levels within the *Nepenthes* radiation (Ornstein-Uhlenbeck shifts)

Gene expression level is a quantitative trait whose phylogenetic evolution can be modelled by a Brownian motion with the additional parameter of an ‘optimum’ value towards which the trait value is drawn, i.e. an Ornstein-Uhlenbeck (OU) process [102]. We tested for diversifying selection on each homolog clusters expression level in this phylogenetic comparative framework, similar to the approach by [21]. Using the R package ouch [102] and our *Nepenthes* species tree (Figure 1), we first fit an OU model with a single expression level optimum (representing stabilising selection). We subsequently fitted nine additional OU models with two expression level optima each which contained a single shift between optima placed at a single internal node of the species tree (diversifying selection, no backward-shifts). Fits of two-optima models were compared against the single-optimum fit using likelihood ratio tests. If after FDR correction [103] any of the two-optima fits was better than the single-optimum fit we concluded that the homolog cluster showed a signature of diversifying selection on its expression level. Terminal nodes (single species) were not tested for separate optima to reduce any possible effect of non-genetic variation (plasticity, technical artefacts) in the selection test.

### Phylogenetically independent contrasts (PICs) of homolog cluster expression levels

As a parameter-free indicator of interspecific variation in gene expression level we chose the mean over the absolute phylogenetically independent contrasts (PICs) per gene (homolog cluster). For this TMM-normalised TPM read counts were used, and averaged over the replicates for *N. rafflesiana* t.f. and *N. hemsleyana*, or the single estimate for the remaining species. PICs [104] can be used to compare quantitative traits between species and are a method to remove possible non-independence of trait observations that might result from shared ancestry and neutral trait evolution (Brownian motion). PICs were calculated using the species tree (Figure 1) and the R package ape [105], for TMM-normalised TPM read counts. Expression levels were averaged over the replicates for *N. rafflesiana* t.f. and *N. hemsleyana*,while we took the single estimate for the remaining non-replicated species.

### Differential gene expression tests

Raw sequence reads of *N. rafflesiana* t.f. and *N. hemsleyana* were mapped against the raw, unfiltered Trinity transcriptome assembly for *N. rafflesiana* t.f. using BWA-MEM [99] with default settings. We did not quality-filter reads [106] or mappings, i.e. multiple placements of reads were tolerated, and all taxonomic and technical ambiguities were retained. Mapped read counts per contig were extracted from .bam files with samtools idxstats, and here we removed contigs that did not contain predicted ORFs, or whose ORFs were not annotating with Embryophyte taxonomy in TrEMBL (see above).

We tested differential gene expression using edgeR [101]. The following comparisons were conducted: (1) fed–control pitchers in *N. rafflesiana* t.f., using three different genotypes each subjected to both treatments, tested in a glm with paired design; (2) pitcher–leaf comparison in *N. rafflesiana* t.f., samples and test design as before; (3) interspecific contrasts of the fed pitchers and leaves of *N. hemsleyana* and *N. rafflesiana* t.f., using first an interactive model (expression level ~ species + tissue + species*tissue), and then separate constrasts within each tissue. Contigs with cumulatively less than ten mappings per million and presence in less than three samples were excluded prior to testing, and counts were TMM-normalised. We used the tag-wise estimate of dispersion, and DGE was considered significant at FDR = 0.05 [103].

### Tests for selection on coding sequence

We tested for the signal of positive selection in each gene based on the ratio of non-synonymous to synoymous substitutions in a codon alignment (d_N_/d_S_ or ω). Purifying (conservative) selection should leave a signal of ω < 1, neutral evolution should produce ω = 1, and positive (diversifying) selection should generate ω > 1. We employed “branch-site” tests, which ask whether for a given branch in a phylogenetic tree, the codons (sites) of a given protein are better classified into only a single ω category or rather several different ω categories including a category of ω > 1 (positive selection). However, we did not want to restrict our molecular evolution hypotheses to specific branches of the *Nepenthes* phylogeny. To get the overall picture, we conducted “branch-site” model tests for all branches in the phylogeny and corrected for multiple testing [107]. We followed this multiple testing procedure in an optimised implementation called Adaptive Branch-Site Random Effects Likelihood (aBSREL) avaliable in the HyPhy software package [24].

After exclusion of *Dionaea*, each homolog cluster within *Nepenthes* was aligned on the peptide sequence in MAFFT v7 [81] and the DNA alignment reconstituted. Alignments contained at least four sequences and 90% presence per codon. Because of the sensitivity of the branch-site test to alignment errors, we evaluated results by posterior filtering on the estimated length of branches yielding significant aBSREL tests, we rejected positive tests when the estimated branch length exceeded 0.1 expected substitutions – it is not conceivable that more substitutions could have accumulated during the relatively young *Nepenthes* crown. Furthermore, tests of molecular evolution can be sensitive to violations of the background species tree [108], hence aBSREL results were conducted with background trees specific to each gene, i.e. estimated from their alignment.

### Testing predictors of signatures of selection

Gene categories were tested against the incidence of signatures of diversifying selection in the radiation of *Nepenthes* (branch-site tests, shifted expression level optima in Ornstein-Uhlenbeck models) using contingency tables (χ^2^-tests) as implemented in the function stats.chi2_contingency from the Scipy package (www.scipy.org). For the quantitative variable ‘mean of absolute PICs of expression levels’, we used permutation tests (R package perm, 100.000 permutations) to test the significance of difference between the means of distributions.

### Gene Ontology enrichment tests

Predicted transcriptome proteins were annotated with GO terms based on their nearest hit in Uniprot SwissProt (blastp, threshold evalue <= 1e-5). We tested GO term overrepresentation in contigs of interest versus background contigs based on simple presence–absence using Fisher’s exact test in the topGO package in R Bioconductor [109]. The “gene universe” was specific for each test and was exactly identical to the set of GO-annotated contigs within which contigs of interest were discovered (i.e. all contigs tested for DGE, or all branch-site tested orthologs). In one enrichment analysis – differentially expressed genes between fed and control pitchers – the superficially dominant gene functions in “fed” pitchers were all related to protein biosynthesis. We thus repeated enrichment tests after masking all contigs annotated with any GO term retrieved by the search for “translation”, “ribosom” and “peptide synth” on http://geneontology.org/ [accessed on 3rd March 2017]. This strategy uncovered previously hidden but neverthless highly significantly enriched GO terms during feeding.

## Acknowledgements

We thank the Ministry of Industry and Primary Resources of Brunei Darussalam for permission to export plant material, and I. and S. Hartmeyer, M. Zehnder and U. Zimmermann for donation of plant material. Claudia Michel helped with the wet lab, and the Genetic Diversity Center Zürich provided computational support. The Functional Genomics Center Zurich is thanked for sequencing, and P. Hunziker for proteomics analyses.

## Author Contributions

MS conducted the experiments, the bioinformatics and statistical analyses, and some of the wet lab procedures; FM and TUG contributed logistic support and materials; MS and AW conceived and designed the study, and wrote the manuscript.

